# Discovering fragment length signatures of circulating tumor DNA using Non-negative Matrix Factorization

**DOI:** 10.1101/2021.06.09.447533

**Authors:** Gabriel Renaud, Maibritt Nørgaard, Johan Lindberg, Henrik Grönberg, Bram De Laere, Jørgen Bjerggaard Jensen, Michael Borre, Claus Lindbjerg Andersen, Karina Dalsgaard Sørensen, Lasse Maretty, Søren Besenbacher

## Abstract

Sequencing of cell-free DNA (cfDNA) is currently being used to detect cancer by searching both for mutational and non-mutational alterations. Recent work has shown that the length distribution of cfDNA fragments from a cancer patient can inform tumor load and type. Here, we propose non-negative matrix factorization (NMF) of fragment length distributions as a novel method to study fragment lengths in cfDNA samples.

Using shallow whole-genome sequencing (sWGS) of cfDNA from a cohort of patients with metastatic castration-resistant prostate cancer (mCRPC), we demonstrate how NMF accurately infers the true tumor fragment length distribution as an NMF component - and that the sample weights of this component correlate with ctDNA levels (r=0.75). We further demonstrate how using several NMF components enables accurate cancer detection on data from various early stage cancers (AUC=0.96). Finally, we show that NMF, when applied across genomic regions, can be used to discover fragment length signatures associated with open chromatin.

## Background

Circulating cell-free DNA (cfDNA) is rapidly emerging as an important biomarker - most notably in cancer and pregnancy. In the cancer setting, the detection of tumor cell derived DNA fragments containing somatically acquired mutations can reveal the presence of cancer. Most approaches rely on the detection of mutations using deep, targeted sequencing of a few genomic regions known to harbor driver mutations for the cancer type of interest[1]. While deleterious or activating mutations in known driver genes are highly specific for cancer, the sensitivity of this approach is constrained as the cancer may not contain the mutation - or the mutations may not be detectable in the blood sample due to low concentration of circulating tumor DNA (ctDNA)[2].

It is, however, possible to get more information out of cfDNA data than just genetic variants. A key difference between cfDNA data and ordinary sequence data is that cfDNA is fragmented *in vivo* by a combination of enzymatic and non-enzymatic processes. Most importantly, during apoptosis, DNA is enzymatically cut between nucleosomes, and hence the lengths and positions of cfDNA fragments reflect the epigenetic state of the cell-types of origin[3,4]. Furthermore, other enzymatic and non-enzymatic fragmentation processes (e.g. oxidative stress) may further contribute to cancer associated fragmentation patterns[5]. In contrast to mutations, these signals are expected to occur across the entire genome and suggest that a focus on sequencing width instead of depth can improve sensitivity.

Indeed, Mouliere *et al.* used exome and shallow whole-genome sequencing (sWGS) to investigate cancer-specific cfDNA fragmentation patterns using human-mouse xenografts or cancer mutations to separate cancer fragments from background cfDNA[6]. They observed a number of cancer-specific distortions - like fragment shortening - and used these to accurately discriminate cancer patients from healthy controls. In their *DELFI* approach, Cristiano *et al.* added a genomic dimension to the analyses by computing the ratio of short (100-150bp) to long (151-220bp) fragments in 5MB windows along the genome and showed how these capture nucleosomal distances, which in turn reflect chromatin state[7]. Using these fragment length profiles as inputs to a machine learning classifier enabled accurate discrimination of earlier stage cancers from controls.

In this manuscript, we introduce a new computational method based on non-negative matrix factorization (NMF), an unsupervised learning method, for simultaneously determining the contributions of different cfDNA sources (e.g. background and tumor) to a sample along with the fragment length signatures of each source[8]. The method is completely unsupervised and uses only fragment length histograms as input and hence provides estimates that are independent of genomic alterations (e.g. SNV, CNV) and sample information like disease state.

## Results

### Discovering tumor fragment length signatures using non-negative matrix factorization

Our approach begins by computing cfDNA fragment length histograms in a series of samples based on paired-end read alignments and then uses these to construct a matrix with cfDNA fragment counts such that each row corresponds to a sample and each column to a specific fragment length (see Fig. 1 for a schematic representation of the workflow). We then normalize the rows of this matrix such that they sum to one before performing NMF, where the input matrix is approximated as the product of two non-negative matrices - both smaller than the input. One of these matrices, the *signature* matrix, has as many columns as the original matrix and represents the preference of observing each fragment length for each cfDNA source. The other matrix, the *weight* matrix, has as many rows as the input matrix and represents the contributions of each cfDNA source to each sample. The number of cfDNA sources is a hyperparameter that needs to be set in advance.

**Fig. 1.**
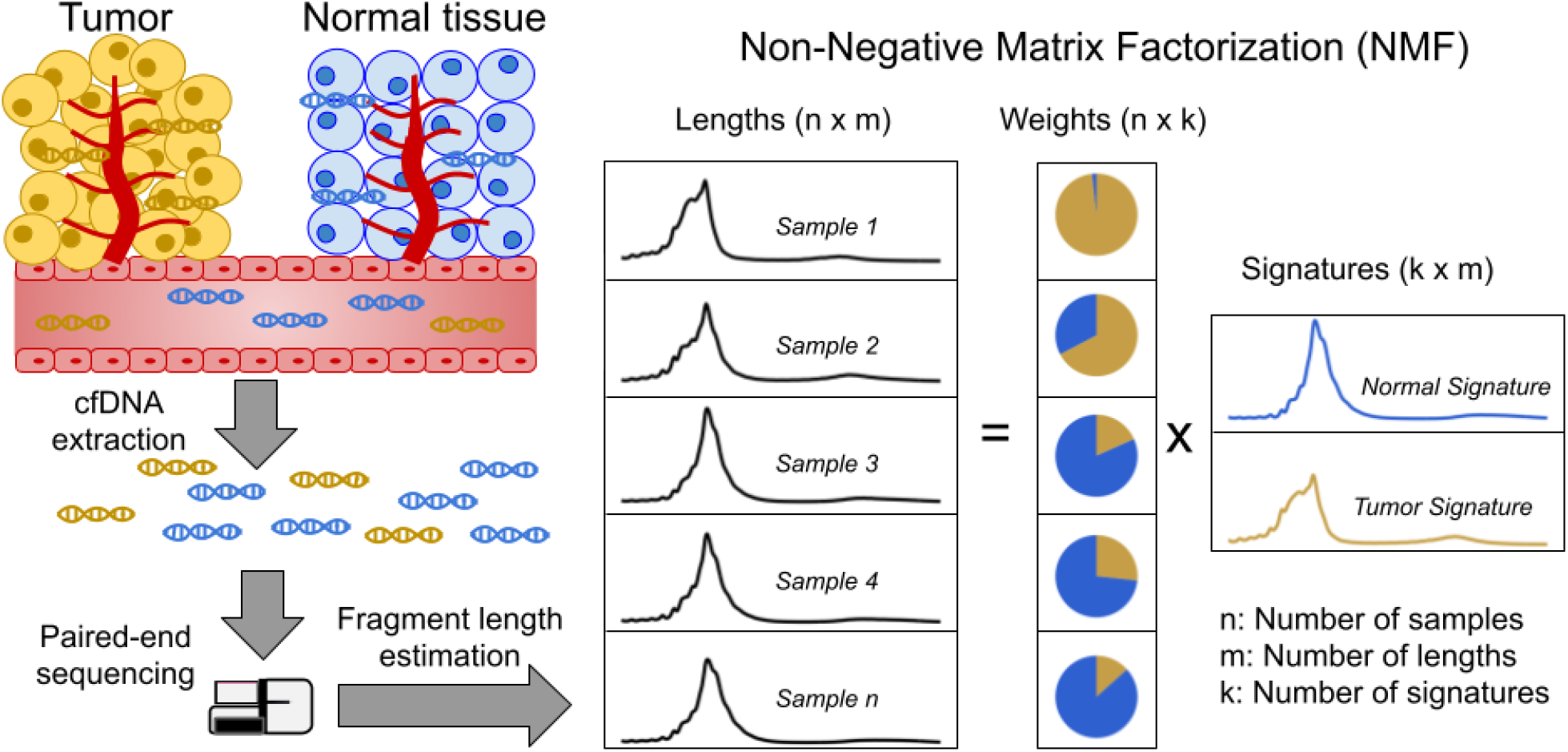
Discovering fragment length signatures using non-negative matrix factorization. The cell-free DNA (cfDNA) pool contains a mixture of fragments from different sources such as tumor cells and background (mainly cells of hematopoietic origin). After performing paired-end sequencing of cfDNA, we estimate fragment length histograms for each sample by aligning reads to the reference genome. We next generate a matrix with fragment length frequencies such that rows and columns represent samples and fragment lengths, respectively. After normalizing the rows of this matrix, we then factorize it into two non-negative matrices: 1) The *signature* matrix is aligned with columns and expresses the preference of each cfDNA source for different fragment lengths and 2) the *weight* matrix, which is aligned with rows, and contains the estimated contributions of each source to each sample.

We first tested our method on shallow whole-genome sequencing (sWGS) of cfDNA (coverage mean: 0.60X, range: 0.36X-0.93X) in 142 plasma samples from 94 patients with metastatic castration resistant prostate cancer (mCRPC). The observed fragment length distributions of high and low ctDNA samples differed (Fig. 2a) and resembled those of previous reports (e.g. Mouliere *et al.*)[6]. Assuming two cfDNA sources (tumor / non-tumor), we then estimated fragment length signatures and weights using NMF on the normalized table of fragment length frequencies. One of the signatures (signature#2) recapitulates key features of the tumor signature previously estimated by Mouliere *et al.* based on mutations including left skew, increased 10 bp periodicity left of the main mode and an enlarged second peak suggesting that this source represents the tumor (Fig. 2b). To confirm the ability of NMF to separate tumor and background (i.e. non-tumor) sources, we performed targeted, deep sequencing (coverage mean: 647X, range:152X-1198X) for a subset of 86 mCRPC patients using a panel of genes related to prostate cancer (5,137 regions, ~1,2 MB)[9,10]. We then called somatic variants in this data and estimated NMF using only fragments that overlap a mutated position. The putative cancer signature could then be compared to the fragment length distribution of fragments with and without mutations (Fig. 2c). The fragment length distribution of fragments containing mutations closely matched the suspected tumor signature estimated using NMF and hence confirms that our method is able to separate tumor and background cfDNA sources solely based on fragment length information.

**Fig. 2.**
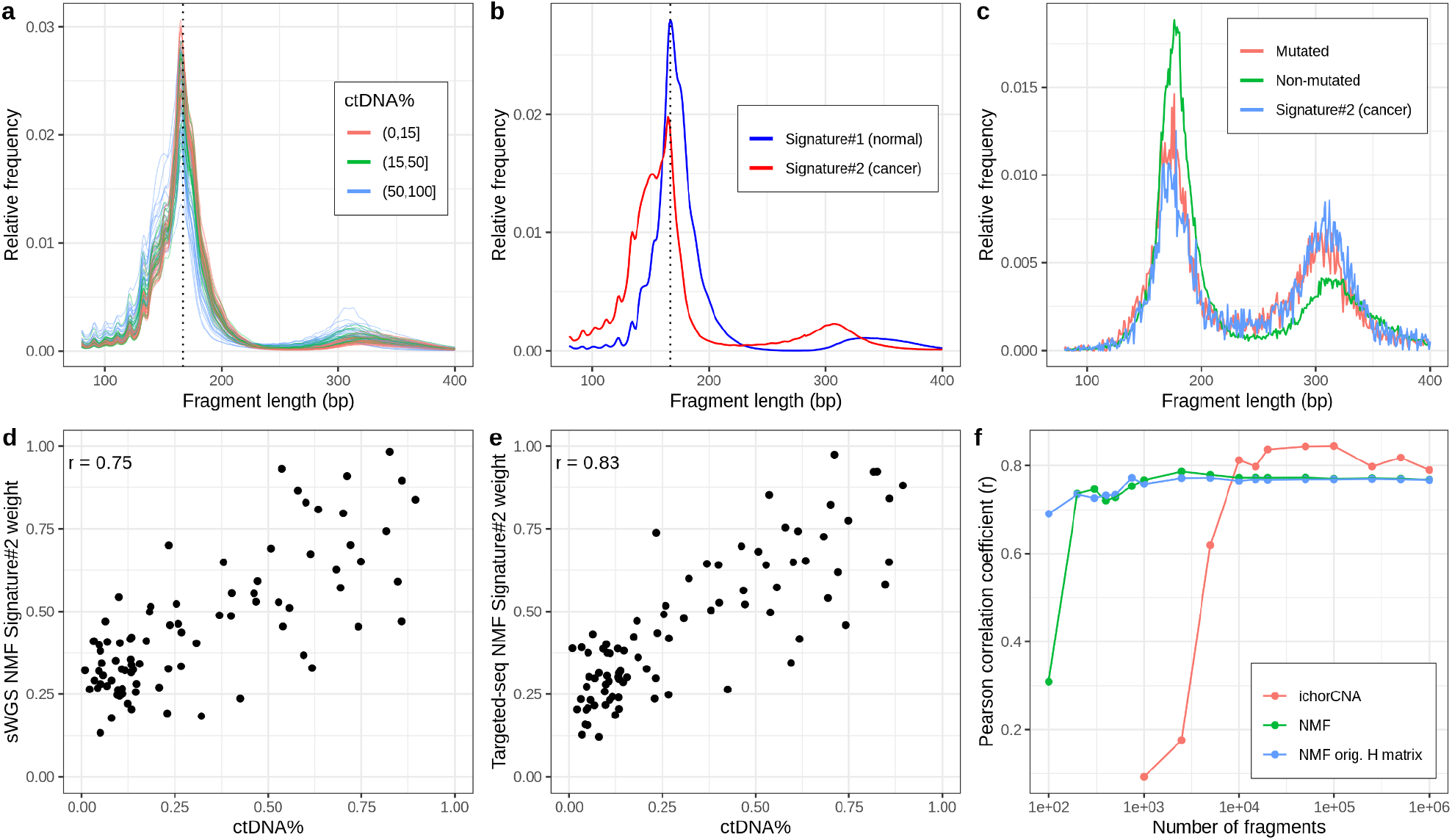
NMF on sWGS and deep targeted sequencing of cfDNA from prostate cancer patients. **a** sWGS fragment length histograms for 86 prostate cancer patients; colors reflect ctDNA fractions estimated from driver variant allele fractions obtained from targeted sequencing performed on the same samples. **b** Fragment length signatures inferred using NMF with two components on the sWGS dataset. **c** Lengths of fragments containing a driver mutation (red dots), lengths of fragments overlapping the mutated position but not containing the mutation (green line) and tumor fragment length signature estimated by NMF (blue line). **d** ctDNA fractions estimated using driver allele frequencies from targeted data versus weights of the second NMF component estimated on sWGS data (signature#2 in panel b). **e** ctDNA fractions estimated using driver allele frequencies from targeted data versus weights of the second NMF component estimated on the same targeted data. **f** Correlation of tumor signature weights estimated using NMF and ichorCNA with ctDNA fractions for different levels of downsampling of the sWGS data.

We next sought to investigate whether the estimated tumor signature weights are related to the blood ctDNA fraction. We, therefore, compared the sample weights of the tumor fragment length signature against sample ctDNA fractions estimated based on driver variant allele fractions (VAFs) obtained from the targeted sequencing data. The estimated weights correlate strongly with ctDNA fraction for both sWGS (r=0.75, Fig. 2d) and targeted sequencing data (r=0.83, Fig. 2e) and hence both confirm the tumor origin of the signatures and demonstrate that our method can be used to estimate ctDNA fraction completely independent of any variant (i.e. SNVs/indels/CNVs) information. Indeed, the estimates obtained using our method were also closely correlated with those determined based on CNV signals using ichorCNA[11] (r=0.70, Supplementary Fig. 1).

We further wanted to assess the relative performance of our variant-independent estimates with those obtained using ichorCNA based on CNVs. On the full dataset, ichorCNA outperformed our method (r=0.79, Supplementary Fig. 2). Yet we speculated that our method might work better when the data is sparse as it leverages information across the entire genome rather than only regions affected by CNVs. We thus ran both NMF and ichorCNA on the sWGS data for different levels of subsampling (Fig. 2f). We observed that, while ichorCNA correlated better with the driver VAF-based ctDNA fractions for higher depths, our method seemed markedly better at low depths. Intriguingly, our method attained correlations greater than 0.68 with the VAF-based estimates using as little as 1000 fragments per sample.

Finally, we also explored using more cfDNA sources in the NMF. More specifically, we ran NMF for up to four components and empirically tested adding the weights of different combinations of NMF components (i.e. assuming different tumor sources) by comparing them with the driver VAF-based estimates (Supplementary Fig. 3). Only marginal improvements were observed using the more complex models suggesting the simpler alternative with two cfDNA sources is preferable as it is more interpretable and leaves little room for overfitting. In resumé, our method is able to determine fragment length signatures and ctDNA fractions on both sWGS and panel sequencing data in a completely unsupervised manner.

### Detecting cancer using multiple fragment length signatures

To see if fragment length signatures can detect the presence of cancer fragments from different cancer types, we reanalyzed the data from the DELFI study[7]. We obtained the raw sWGS data (mean coverage: 2.84X range: 0.71X-13.4X) from 498 samples from this study including 260 healthy controls and 238 cancers distributed across seven different cancer types. The fragment length distributions in these samples (see Fig. 3a) were similar to the prostate cancer data, and as expected, we saw a general tendency for shorter fragment lengths in cases compared to controls. There were, however, also visible differences between the prostate data and the DELFI data set. We observed, for instance, fewer di-nucleosome fragments in both DELFI cases and controls compared to the prostate cancer data. Like the prostate analysis, we trained NMF on the matrix containing fragment length histograms for each sample, again assuming two cfDNA sources (see Fig. 3b). We assumed that the signature with the lower mean fragment length was the cancer-related signature. This signature did indeed tend to have a higher weight in the cancer samples (Supplementary Fig. 4a), and it could differentiate between cancer and control samples with an AUC of 0.75 (Supplementary Fig. 4b). We then tested whether using more signatures would improve this classification. We used a linear Support Vector Machine (SVM) to see if we could separate cancer and control samples based on their signature weights for a given number of signatures. The results showed that adding more signatures significantly improved the AUC (see Fig. 3c). The classification continued to get better until we reached ~30 signatures, and for 30+ signatures, we got an AUC above 0.95. This AUC is comparable to the AUC of 0.94 the DELFI method achieves by using Gradient Boosting Machine on ~500 features per sample.

**Fig. 3.**
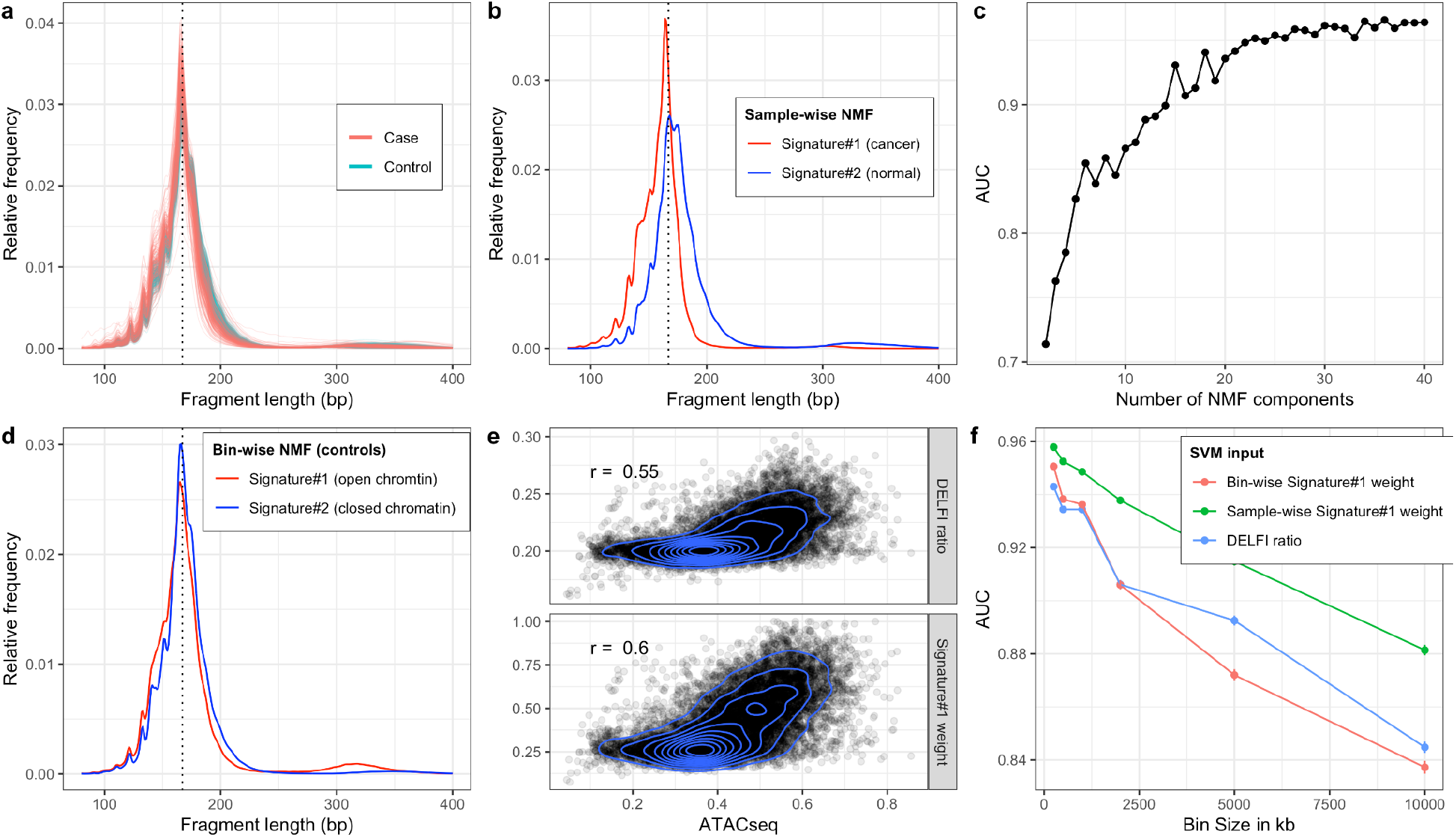
NMF on cfDNA sWGS from the DELFI study. **a** sWGS fragment length histograms for the 533 DELFI samples; colors indicate case-control status of the sample. **b** Fragment length signatures inferred using NMF with two components on the sWGS dataset. **c** AUCs obtained when discriminating cases versus controls using a linear SVM on the sample component weights across different numbers of components in the NMF model. **d** Epigenetic fragment length signatures estimated using fragment length histograms from 250kb bins along the genome aggregated across all control samples. **e** Ratio of short (100-150bp) to long (151-220bp) fragments (“DELFI ratio”) or weight of the first NMF component (signature#1 in panel d) versus ENCODE ATACseq from a Lymphoblastoid cell-line for 250kb genomic bins. **f** AUCs obtained when discriminating cases versus controls using a linear SVM on “DELFI ratio” or weight of the first NMF components from panel d (blue) or panel b (green) inferred in bins along the genome for different bin sizes.

### Estimating epigenetic fragment length signatures using NMF

The DELFI method also uses fragment length information to detect cancer samples, but rather than looking at the total length distribution, it looks at fragment lengths in bins along the genome[7]. The DELFI creators have shown that the length distribution of cfDNA fragments in a genomic region carries information about its epigenetic state, and that such information can be used to distinguish cancer samples from healthy controls. Specifically, the DELFI method uses the ratio of short (100-150bp) to long (151-220bp) fragments in 5MB windows along the genome as input to a machine learning classifier. We wished to investigate whether we could use NMF fragment length signatures to better capture this epigenetic state information and hence yield better classification. First, we partitioned the genome into non-overlapping bins of 250kb and computed the fragment length histograms for each bin in each sample. We then summed histograms for each bin across all healthy controls to yield a matrix with rows corresponding to genomic bins instead of samples as before, and ran NMF using two signatures. This resulted in the two *bin-wise* NMF signatures shown in Fig 3d. To compare, we also calculated the short-to-long ratio used by the DELFI method in each bin. Looking at chromatin accessibility in a lymphoblastoid cell-line measured by ATACseq for the 250kb bins, we observed a slightly better correlation for the weight of the first signature than the DELFI ratio (Fig 3e). We then estimated the first bin-wise signature’s weight in each bin in each sample and used these weights as input to an SVM and assessed classification performance across different bin sizes using the two inputs. The results in Fig 3f show that for small bin sizes, we get a slightly better result by using the bin-wise NMF weight for each bin instead of the short-to-long DELFI ratio as SVM input. But for larger bin sizes, the Delfi ratio is better than using the bin-wise NMF signature. Finally, we also tried inferring the weight of the sample-wise NMF signature from Fig3b for each bin and using that as input to the SVM (green line in Fig3f). This turned out to give superior results compared to using the bin-wise NMF signature.

## Discussion

In this article, we propose non-negative matrix factorization (NMF) of fragment length distributions as a new tool in the cfDNA-seq analysis toolbox. A key objective in cfDNA-seq analyses in the cancer setting is to determine the amount of circulating tumor DNA - most importantly whether any tumor DNA is present at all. This carries immense importance both for early detection of cancer in the screening setting - and for detection of relapse after treatment, due to residual disease after surgery or resistance to chemotherapy.

Most analyses have focused on using mutational signals for determining ctDNA load through estimation of variant allele fractions for SNVs in deep, targeted cfDNA-seq data[1] - or CNVs by sWGS[11]. Yet non-mutational signals such as those shown by several studies to be manifested in the fragment length distribution are gaining traction as they may improve sensitivity by enabling aggregation of the cancer signal across the entire genome rather than just positions affected by mutations. So far, fragment length signals have been approached either using a simple summary statistic like the ratio of short to long fragments (e.g. DELFI[7]) or by manually curating features reflecting the distribution and then use these as input for a supervised machine learning algorithm (e.g. Mouliere *et al.[6]*). However, manual featurization may not make the best use of all relevant information contained in the distribution - and supervised learning with many features carries the risk of overfitting.

We propose NMF as a general analytical framework for working with cfDNA-seq fragment length distributions. NMF enables us to simultaneously estimate fragment length *signatures* and their *weights* in each sample. Using sWGS cfDNA-seq data from a cohort of patients with metastatic prostate cancer, we show that NMF estimated with two components discovers a signature, which accurately matches the true tumor fragment length distribution and exhibits many of the characteristics previously associated with ctDNA. The weights of this signature correlated strongly with ctDNA levels - nearly as good as ichorCNA - without using any information about variants or ctDNA levels. Importantly, similar results were obtained when using deep, targeted cfDNA-seq. Furthermore, subsampling experiments revealed that NMF was markedly more robust when less data is available than ichorCNA; we speculate this also implies that NMF works better at low tumor fractions although this assertion could not be directly tested on the prostate cancer data. Finally, as the fragment length signal is likely orthogonal to any mutational signal, it may be possible to combine these lines of information to obtain a better, joint estimate of tumor load.

The unsupervised nature of NMF implies little risk of overfitting and the ability to inspect the fragment length profiles of each signature provides full transparency of the method in contrast to many other approaches such as the supervised learning by Random Forest strategy applied by Mouliere *et al.* for determining ctDNA levels. Transparency is important because using non-mutational information carries the risk of using information that is not directly linked to the presence of ctDNA in blood, but instead reflects e.g. an ongoing immune response. This in turn may impact a models’ ability to generalize to unseen data and clinicians’ trust in the model. Using NMF, we were able to verify that the fragment length profile of the signature correlating with ctDNA levels does indeed match the length distribution of fragments containing mutations and hence that the model is directly measuring ctDNA load. Hence, the combination of unsupervised learning and transparency suggest that NMF constitutes a robust modelling framework for cfDNA-seq length spectra.

The data from the prostate cancer cohort generally contained patients with a high tumor burden and we wished to also test the models applicability in a screening context characterised by low ctDNA load and also look at different cancer types. We obtained access to the data from the DELFI study, which contains cfDNA-seq data from a range of cancer types and primarily from patients with early stage disease[7]. The fragment length distributions from high ctDNA load prostate cancers and those of DELFI cases shared features (e.g. left skew of the main mode), but also differed as for instance the second mode of the distribution was less pronounced in the DELFI data. These differences may reflect differences in sample processing (e.g. DNA extraction method) and sequencing technology rather than actual biological differences between the studies suggesting that transfering models trained on one dataset to another may be difficult although assertion was not directly tested in the present study. Using an unsupervised method such as NMF to learn the relevant fragment length signatures can alleviate this transferability problem. To know which of the two signatures learned corresponds to cancer fragments, one could use a previous set of signatures trained on a labeled set as the starting point for the NMF optimization.

On the DELFI dataset we furthermore demonstrated that using several NMF components enabled accurate cancer detection of early stage cancers - on par with the original DELFI results. We obtained the best classification results by using 30 or more signatures, and an even larger number of signatures could likely be relevant for larger or more heterogeneous datasets. Using a model with tens of parameters rather than hundreds as in the genomic window model makes overfitting less likely.

Finally, we investigated whether the NMF approach could improve upon the genomic bin-based approach proposed by Cristiano *et al.* We first showed how NMF can discover fragment length signatures of different chromatin states when trained across genomic bins, where the fragment length histogram in each bin has been aggregated across multiple samples (*bin-wise* training). The learned epigenetic signatures turned out to correlate better with open chromatin as measured using ATACseq than the DELFI ratio, but did not yield a clear improvement in classification performance over the DELFI method. We speculated that the lack of classification improvement could be due changes between cases and controls not related to chromatin status. We therefore investigated whether the bin-based approach could be improved by instead inferring the signature weights of the *sample-wise* trained NMF model in each genomic bin. To our surprise, this model outperformed both the bin-wise NMF and the DELFI ratio across all bins sizes, which may indicate that the DELFI classification signal is not purely an epigenetic signal but in part caused by CNVs in the tumors or other cancer-specific distorsions.

## Conclusions

In resumé, we here demonstrate the use of NMF as a general and robust statistical approach for analysing fragment length distributions from cfDNA-seq.

## Declarations

### Ethics approval and consent to participate

The prostate study was approved by The National Committee on Health Research Ethics (#1901101) and notified to The Danish Data Protection Agency (#1-16-02-366-15). All patients provided written informed consent.

### Consent for publication

Not applicable.

### Availability of data and materials

The prostate cancer datasets generated during the current study are available from the corresponding authors upon reasonable request and pending further ethical approval and data sharing agreements, as required by Danish law. The raw DELFI data are available through the European Genome-phenome Archive (EGA accession: EGAD00001005339). Any data obtained in this study as a derivative of the DELFI data is available upon request pending approval by the responsible EGA Data Access Committee and permission from the original study authors. Software for calculating fragment lengths and applying NMF is available under the GPL available at: http://grenaud.github.io/cinch/. All other code used to perform analyses in this study is available upon reasonable request.

### Authors’ contributions

LM and SB conceived the idea of applying NMF to cfDNA fragment length data. GR, MN, LM and SB analyzed the data. MN, JL, HG, BDL and KDS designed and/or performed sWGS and targeted, deep sequencing experiments on the prostate cancer samples. JBJ and MB were responsible for patient recruitment, contributed samples and clinical information. CLA helped interpret the results. GR, LM and SB drafted the manuscript with input from all authors. All authors approved the final manuscript.

## Acknowledgements

The authors wish to thank the staff at the departments of urology at Aarhus University Hospital and Regional Hospital West Jutland for patient recruitment and collection of clinical data. We would also like to thank lab technicians and clinical academics at the Department of Molecular Medicine (AUH, Denmark) and the Department of Medical Epidemiology and Biostatistics (Karolinska Institutet, Sweden) for excellent assistance throughout the project.

## Methods

### Sample processing and DNA extraction

A total of 142 EDTA-blood samples were collected from 94 patients with metastatic castration-resistant prostate cancer at Aarhus University Hospital and Regional Hospital of West Jutland between April 2016 and August 2019. Samples were collected at treatment initiation and/or at disease progression (1^st^ – 4^th^ line treatment). A total of 36 patients had multiple samples (of these: median n=2 samples/patient (range: 2-4 samples/patient)).

Blood samples were collected in BD Vacutainer K_2_ EDTA tubes (Beckton Dickinson) and processed within 2 hours. cfDNA was extracted from 2.0-4.5 mL (median: 4.0 mL) plasma on a QIAsymphony robot (Qiagen) using the QIAamp Circulating Nucleic Acids kit (Qiagen) as described by the manufacturer. cfDNA concentration was determined by droplet digital PCR (ddPCR) using a QX200 AutoDG Droplet Digital PCR System (Bio-Rad) according to the manufacturer’s instructions as previously described [12]. Germline DNA from buffy coats (peripheral blood mononuclear cells (PBMC)) was extracted on a QIAsymphony robot (Qiagen) using the QiaSymphony DSP DNA Mini Kit (Qiagen) following manufacturer’s instructions. DNA concentrations were determined using Qubit fluorometric quantification (Qubit dsDNA Broad range, ThermoFisher). Prior to library preparation, the Covaris E220 Evolution ultrasonicator (Covaris) was used to shear germline DNA to shorter fragments (~250-350 bp).

### Library preparation and shallow whole-genome sequencing

Libraries for next generation sequencing were prepared using the Kapa Hyper Library Preparation Kit (KAPA Biosystems) with 7.0-50.0 ng cfDNA (median 30.5 ng) or 50 ng for germline DNA as input. xGen CS-adapters – Tech Access (IDT-DNA) with Unique Molecular Identifiers (UMIs) on both strands and primers containing unique indexes on both strands were used to generate indexed libraries. Plasma libraries were pooled equimolarly and paired-end sequenced (2×151bp) on an Illumina® Novaseq instrument (S-prime flowcell), generating 12.66-34.30 million read pairs/ sample (median: 19.99) corresponding to coverages from 0.36-0.93X (median: 0.60X). Fastq files were demultiplexed using bcl2fastq (v2.20.0.422) and quality checked using fastQC and fastqScreen (Available from: http://www.bioinformatics.babraham.ac.uk/projects).

### Estimation of fragment lengths

We used leeHom with the “--ancientdna” option on the raw fastq files to strip adapters, and, where possible, to reconstruct DNA fragments by merging overlapping paired-end reads. UMIs were removed by trimming the first five nts of reach read[13]. Reads were then mapped to hg19 using BWA-MEM v.0.7.17 with seed length (“-k”) set to 19[14]. PCR and optical duplicates were removed using samtools rmdup using option “-s” for single-end reads. Paired-end reads with mapping quality below 30, mapping within ENCODE excluded regions (ENCFF001TDO) or which contained soft or hard clips in either of the two reads were filtered away before computing fragment lengths. Fragment lengths were then directly obtained as the length of the reconstructed fragment when reconstruction was possible and otherwise obtained from the insert size calculated by the aligner based on distances on the reference sequence. Finally, fragment lengths less than 30 or greater than 700 were discarded and a matrix with fragment length counts constructed such that rows and columns corresponded to samples and fragment lengths, respectively.

### Non-negative matrix factorization

The rows of the matrix with fragment length counts were first scaled such that they sum to one. Non-negative matrix factorization (NMF) of the normalised matrix was then performed using scikit-learn (sklearn.decomposition.NMF) with random initialization, multiplicative updates and the Kullback-Leibler loss function. For the bin-wise NMF experiments, the NMF was repeated across 20 different random initializations and the fit achieving the lowest loss selected. The estimated fragment length signature and weight matrices were scaled to sum to one for each signature and sample, respectively.

### Analysing fragment lengths in bins along the genome

We first divided the genome into bins of 250kb and estimated the fragment length distributions for each sample in each bin. We then calculated the mean number fragments in each bin across the controls. To avoid bins with possible mapping problems, we excluded bins where the mean number of fragments were less than the median-2*IQR or greater than the median+2*IQR. Comparison with ATACseq data was performed using ENCODE track “ENCFF603BJO” (fold change over control, GM12878 cell line) lifted from hg38 to hg19.

### Building support-vector machine classification models

The classification results based on multiple fragment length signatures and the classification results on epigenetic signatures and DELFI ratios were calculated using linear SVMs implemented in R using tidymodels and kernlab. Before training, the values for each sample was standardized by subtracting the mean and dividing with the standard deviation. Accuracy and AUC were calculated using repeated 10-fold cross-validation (50 repeats).

### Deep targeted prostate-tailored sequencing

A total of 85 indexed libraries were subjected to deep targeted sequencing based on ichorCNA ctDNA% estimates from plasma cfDNA libraries. A previously designed gene panel that captures regions in the human genome commonly altered in PC was used for capture[9,10]. Libraries were pooled equimolarly (8-plex) for in-solution target enrichment using Twist Bioscience’s Custom Target Enrichment. Captured pools were paired-end sequenced (2×100bp) on an Illumina Novaseq instrument (S1 flowcell). Fastq files were generated as for the sWGS data and adaptor trimming, mapping, duplicate removal, realignment and quality score recalibration performed as described above for the ichorCNA workflow for sWGS data. The resulting depth of coverage for the targeted regions was on average 647X with a minimum of 152X and a maximum of 1198X.

### Variant calling and interpretation

Somatic variants were called using VarDict (v. 1.6)[15], Strelka2 Somatic (v. 2.9.10) [16], GATK mutect2 (v. 4.1.2.0)[17], and VarScan2 (v. 2.4.2) [18]. For each sample, a patient-matched germline sample was used as control. The impact of each variant was annotated using the Ensembl Variant Effect Predictor (ensemble-vep v. 96.0) [19]. Somatic variants were required to be supported by at least 10 reads, called by ≥3 callers, and annotated as either pathogenic or likely pathogenic in OncoKB or ClinVar [20,21] or introduce a premature stop or frameshift in the coding sequence. High-impact variants called by 2 callers were not discarded whereas low-frequency variants (VAFs <0.02) were discarded unless they had high impact, were called by all 4 callers, or were detected in another sample from the same individual. LOH-status was annotated for each variant using cfDNA copy number profiles and allele ratio of heterozygous SNPs. All variants were manually curated in IGV (v. 2.5.3).

### Running ichorCNA

Adapter sequences were trimmed using Cutadapt (v1.16)[22] and paired-end sequences mapped to the hg19 reference genome using BWA MEM (v0.7.15-r1140)[14]. PCR and optical duplicates were removed from each library independently using Samblaster (v0.1.24)[23] and the final bam files realigned (GATK v3.8.1.0)[24]. ichorCNA was run with default parameters [11].

### Estimation of ctDNA fractions from targeted seq data

CtDNA fractions were also estimated from the targeted sequencing data. First, the tumor cell purity (i.e. tumor cell fraction) was calculated from somatic mutations with moderate or high impact. In brief, for each sample the somatic mutation with the highest VAF was used (i.e. clear driver mutation). If multiple mutations had similar VAFs (+/− 2%), the median VAF of these mutations were used instead. A total of 1-2 mutations per sample were used to estimate purity. Accounting for LOH status of the mutation(s), the tumor cell purity was estimated as follows: purity=2*VAF (no LOH) or purity=2/(1/VAF+1) (LOH).

Next, to obtain ctDNA fractions, the tumor cell purity was adjusted for tumor ploidy: ctDNA fraction = tumor cell purity * tumor ploidy/(tumor cell purity * tumor ploidy + normal cell purity * normal ploidy). Here, normal purity was set to 1-tumor cell purity and normal ploidy was set to 2. Tumor ploidy estimates were obtained from PureCN[25].

## Supplementary Material

**Supplementary Fig 1.**
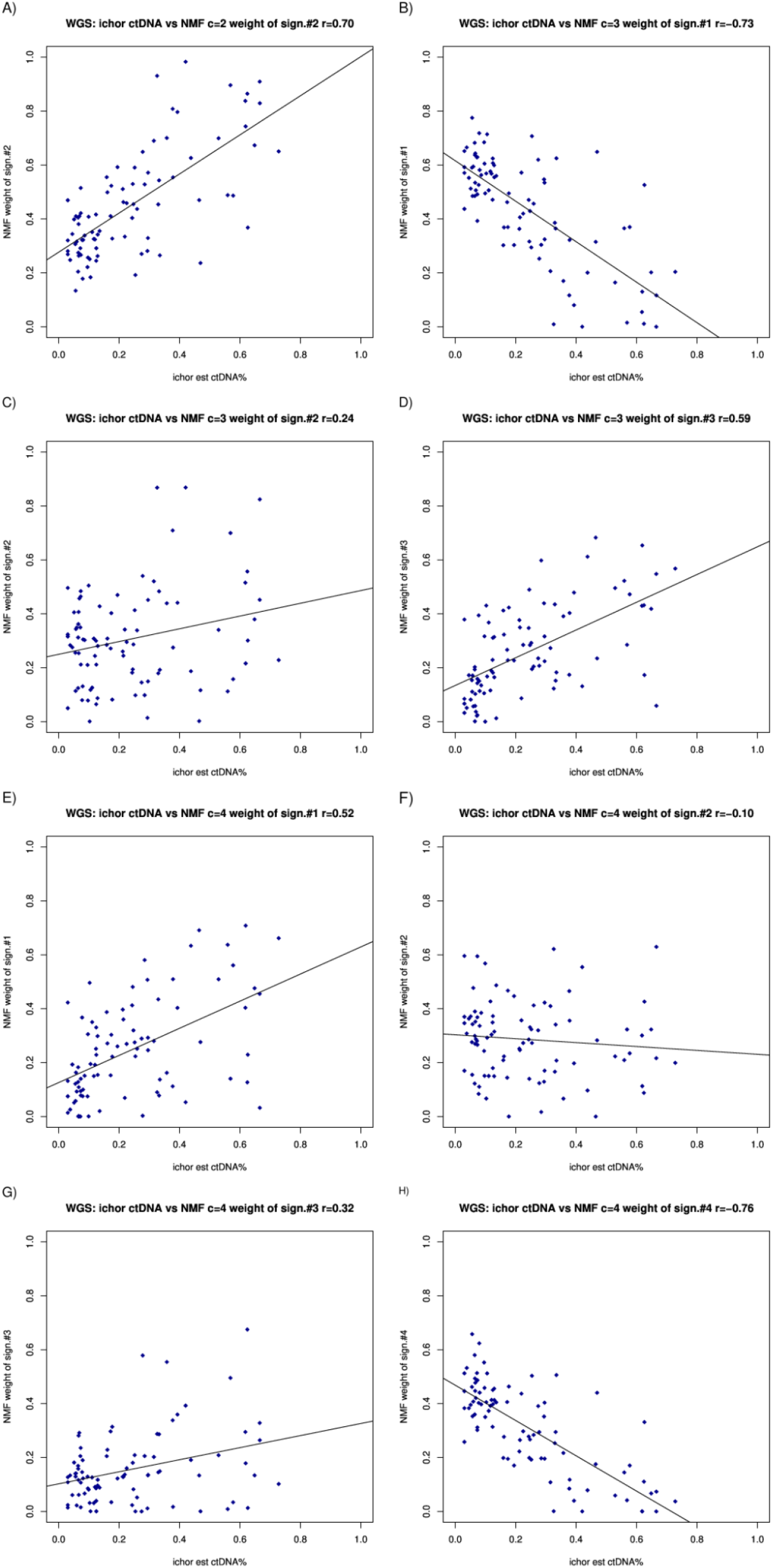
Correlation between the estimates of ichorCNA and NMF components for the whole-genome data.

**Supplementary Fig 2.**
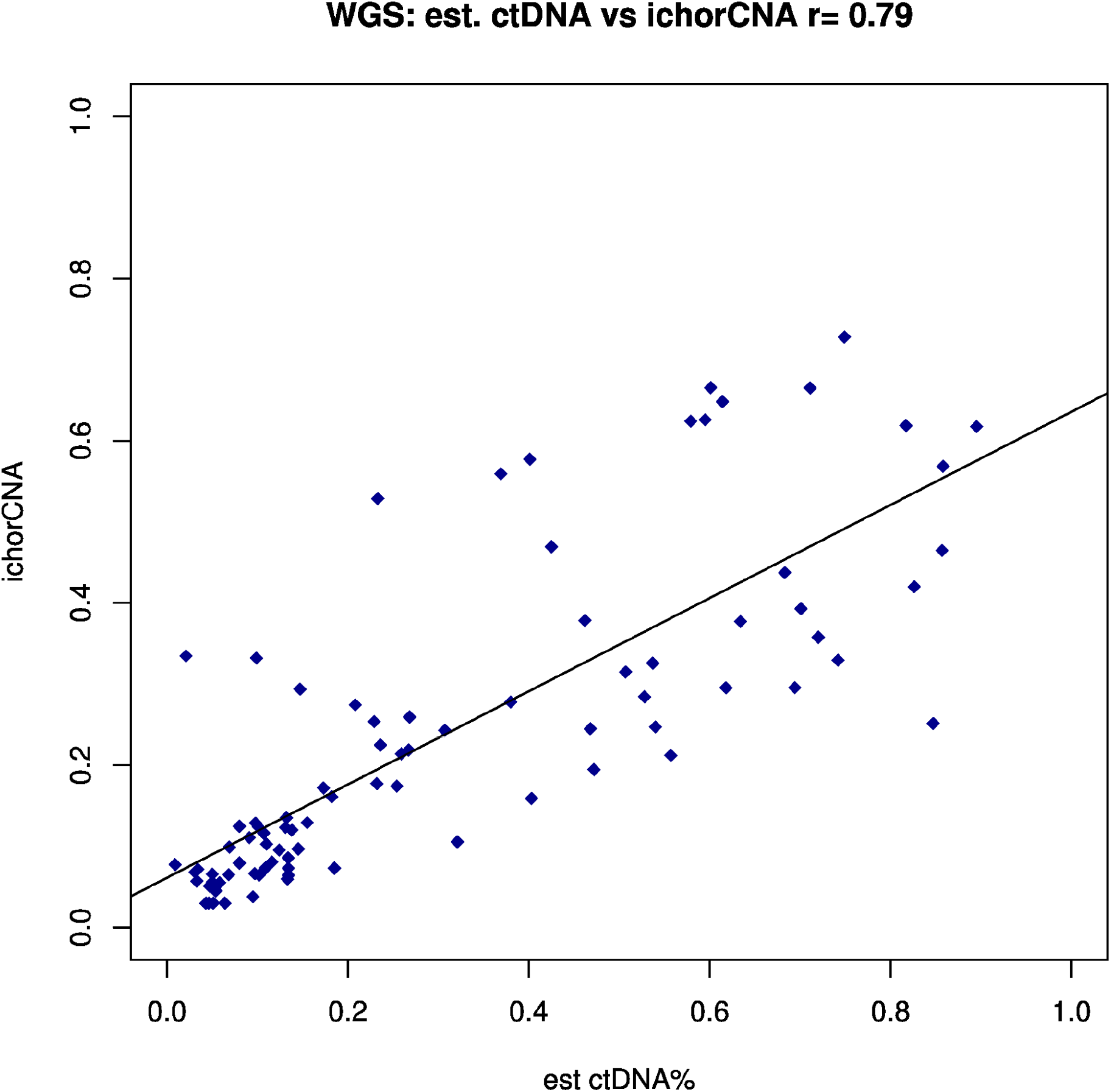
Correlation between predicted ctDNA and ichorCNA for the whole-genome data.

**Supplementary Fig 3.**
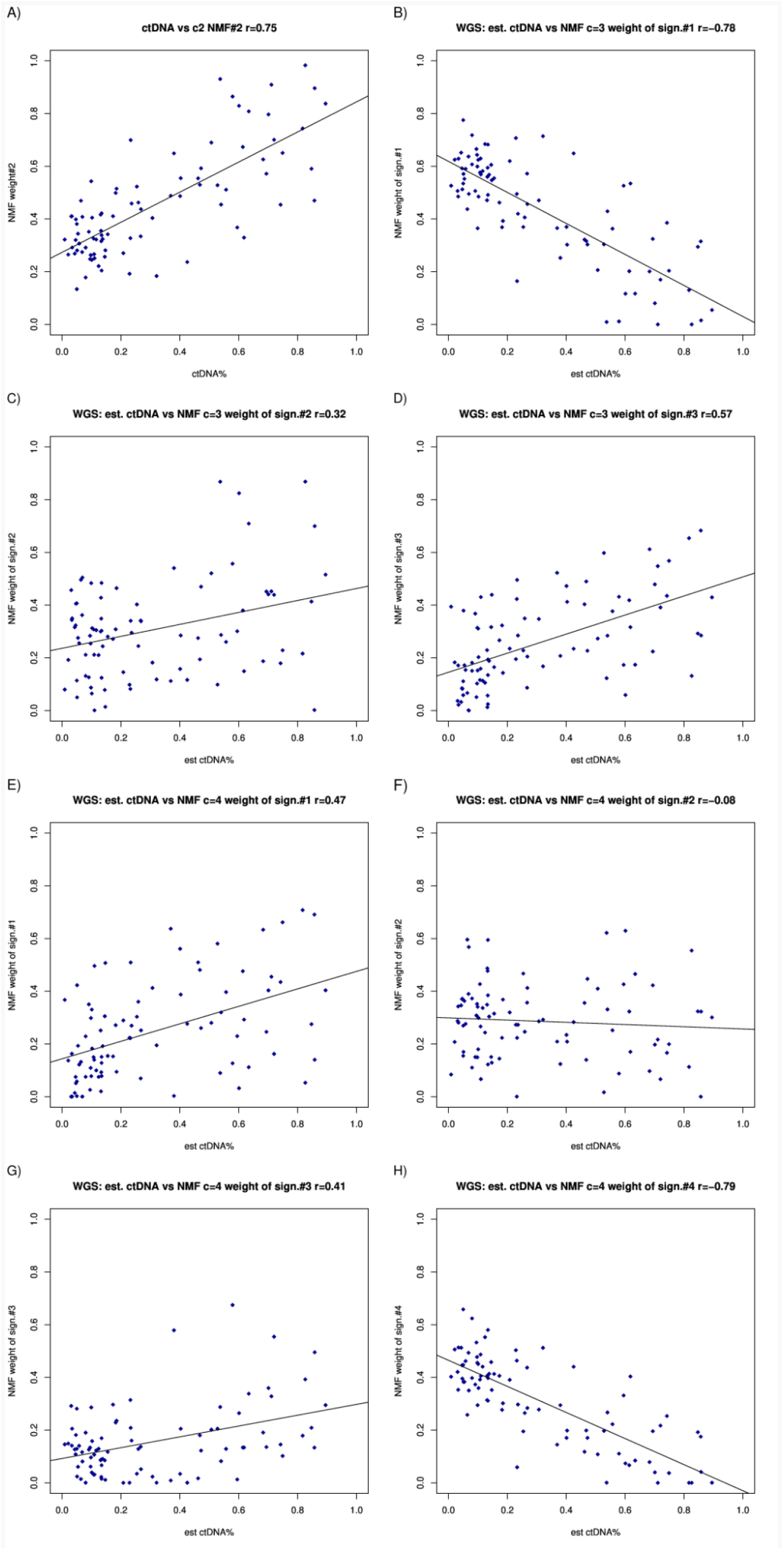
Correlation between estimated ctDNA and NMF components for the whole-genome data.

**Supplementary Fig 4.**
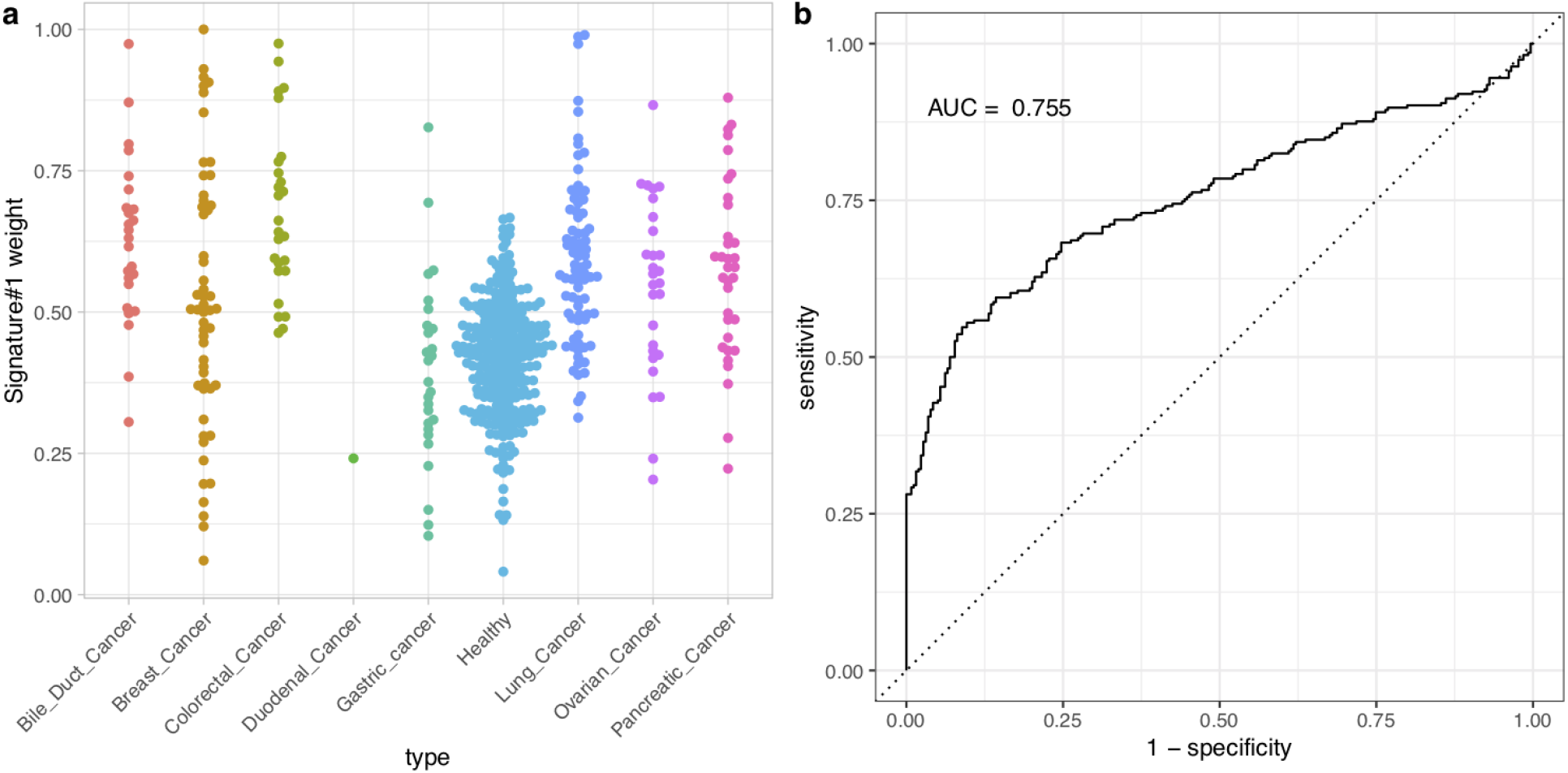
Using NMF with two fragment length signatures to classify sample type on DELFI data. a) The distribution of Signature #1 weights for different sample types. b) ROC curve for cancer vs control classification using Signature #1.

